# Zebrafish Drug Screening Identifies Erlotinib as an Inhibitor of Wnt/β-Catenin Signaling and Self-Renewal in T-cell Acute Lymphoblastic Leukemia

**DOI:** 10.1101/2023.08.28.555200

**Authors:** Majd A. Al-Hamaly, Anna H. Cox, Meghan G. Haney, Wen Zhang, Emma C. Arvin, Shilpa Sampathi, Mary Wimsett, Chunming Liu, Jessica S. Blackburn

## Abstract

The Wnt/β-catenin pathway’s significance in cancer initiation, progression, and stem cell biology underscores its therapeutic potential, yet clinical application of Wnt inhibitors remains limited due to challenges posed by off-target effects and complex crosstalk with other pathways. In this study, we leveraged the zebrafish model to perform a robust and rapid drug screening of 773 FDA-approved compounds to identify Wnt/β-catenin inhibitors with minimal toxicity. Utilizing zebrafish expressing a Wnt reporter, we identified several drugs that suppressed Wnt signaling without compromising zebrafish development. The efficacy of the top hit, Erlotinib, extended to human cells, where it blocked Wnt/β-catenin signaling downstream of the destruction complex. Notably, Erlotinib treatment reduced self-renewal in human T-cell Acute Lymphoblastic Leukemia cells, which are known to rely on active β-catenin signaling for maintenance of leukemia-initiating cells. Erlotinib also reduced leukemia-initiating cell frequency and delayed disease formation in zebrafish models. This study underscores zebrafish’s translational potential in drug discovery and repurposing, and highlights a new use for Erlotinib as a Wnt inhibitor for cancers driven by aberrant Wnt/β-catenin signaling.

**Highlights:**

- Zebrafish-based drug screening offers an inexpensive and robust platform for identifying compounds with high efficacy and low toxicity *in vivo*.
- Erlotinib, an Epidermal Growth Factor Receptor (EGFR) inhibitor, emerged as a potent and promising Wnt inhibitor with effects in both zebrafish and human cell-based Wnt reporter assays.
- The identification of Erlotinib as a Wnt inhibitor underscores the value of repurposed drugs in developing targeted therapies to disrupt cancer stemness and improve clinical outcomes

**Graphical Abstract:** 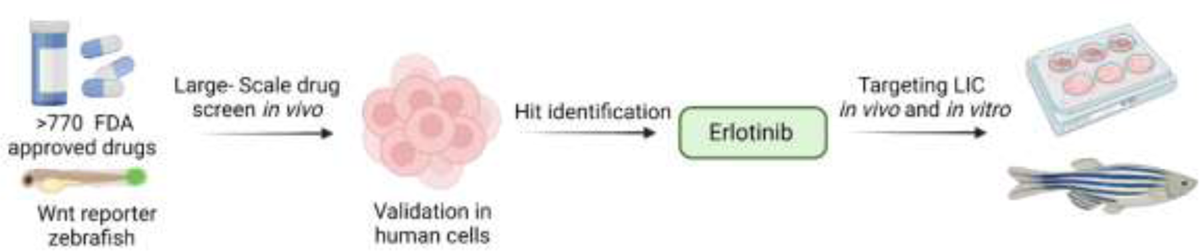

## 1. Introduction

Nearly half of all solid tumors and many leukemias have aberrations in the Wnt signaling cascade [1–3], which is a highly conserved cellular signaling pathway that plays a multifaceted role in both normal cellular processes and cancer development [4, 5]. Canonical Wnt signaling involves the binding of Wnt ligands to cell surface receptors, which subsequently activates a series of intracellular events that culminate in the stabilization and translocation of β-Catenin into the nucleus. Within the nucleus, β-Catenin interacts with T-cell factor/lymphoid enhancer factor (TCF/LEF) transcription factors to activate the transcription of Wnt target genes associated with proliferation, differentiation, and cell fate determination [6].

In the context of cancer, aberrant Wnt/β-Catenin signaling frequently arises due to mutations in key components of the pathway, leading to dysregulated β-Catenin accumulation and unchecked transcriptional activation. Hyperactivation of Wnt/β-Catenin signaling has been linked to the onset and progression of many cancers, including colorectal, breast, hepatic, brain, and blood [7–10]. Varied levels of Wnt/β-Catenin signaling within a tumor also reflect intratumoral heterogeneity, and can account for distinct cellular functions in a tumor, such as proliferation, self-renewal, and epithelial-to-mesenchymal transitions [11].

Importantly, accumulating evidence suggests a pivotal role for this signaling axis in the maintenance and propagation of cancer stem cells (CSCs), a subpopulation of tumor cells endowed with self-renewal and differentiation capabilities similar to normal tissue stem cells [12, 13]. For example, 80% of colorectal cancers have mutations in the adenomatous polyosis coli (APC) gene, leading to loss of APC function and aberrant β-Catenin accumulation in the nucleus, fostering activation genes associated with colorectal cancer stem cell self-renewal and contributing to therapy resistance [7]. Similarly, breast cancer stem cells exploit Wnt signaling for enhanced stem-like traits and tumorigenesis [14]. Increased Wnt signaling in certain subtypes of breast cancer is associated with more aggressive tumor phenotypes and poor patient prognosis [15]. CSCs in both hepatocellular carcinoma and glioblastoma utilize dysregulated β-Catenin signaling to drive stem cell features, including tumorigenesis and therapy evasion [16, 17].

Wnt signaling is also crucial in T-cell Acute Lymphoblastic Leukemia (T-ALL), where activating mutations in interconnected pathways like NOTCH1 result in an upregulation of β-Catenin in ∼85% of patients and subsequent high expression of Wnt target genes including c-MYC, TCF, and LEF [18]. Intriguingly, single cell gene expression analysis of samples from T-ALL patients with minimal residual disease (MRD) revealed a high expression of β-Catenin within the leukemia stem cell niche [19]. Giambra et al. further demonstrated that active Wnt signaling is confined to a subset of cells enriched for leukemic stem cells using a stably integrated Wnt reporter [20]. Thus, the Wnt/β-Catenin signaling axis has emerged as a critical driver of stemness, drug resistance, and tumorigenesis across diverse cancers, emphasizing its potential as a therapeutic target.

Wnt inhibitors, despite their potential therapeutic promise, have yet to find widespread clinical application due to several challenges. The involvement of Wnt signaling in normal tissue functions, such as embryonic development, cellular homeostasis, and tissue regeneration, raises concerns about detrimental side effects upon pathway inhibition [21]. Furthermore, the multi-faceted crosstalk between Wnt signaling and other crucial pathways necessitates a thorough understanding of potential off-target effects [22, 23]. While novel molecules for Wnt inhibition have been identified [24, 25], the lengthy and costly drug development process, along with the need for rigorous pre-clinical and clinical evaluations, has hindered their swift translation into clinical practice. These challenges highlight the importance of finely tuned strategies to harness the therapeutic potential of Wnt/β-Catenin inhibition while mitigating associated risks.

Zebrafish drug screening presents a compelling avenue for identifying novel Wnt inhibitors with minimal off-target effects. The genetic similarities between zebrafish and humans, with approximately 80% shared disease-associated proteins [26], underscore the translational relevance of findings made in zebrafish. This similarity, coupled with the growing body of evidence on conserved pharmacology, metabolism, and drug targets [27, 28], lends confidence to the predictive value of zebrafish-based results. The ability to generate a significant number of embryos weekly enables large-scale screenings, while automated methodologies for treatment administration, imaging, and analysis streamline the process and enhance reproducibility [29–31]. The optical transparency of zebrafish larvae also facilitates real-time monitoring of drug responses and their potential side effects, offering insights into drug impact with high spatial and temporal resolution [32]. In total, zebrafish-based drug screens afford a holistic understanding of drug effects, which can enable the identification of promising Wnt inhibitors while minimizing off-target effects, potentially hastening the translation of discoveries into the cancer clinic.

Here, we utilized a transgenic strain of zebrafish, in which cells with activated Wnt/β-Catenin signaling are GFP-positive, to screen a library of 773 FDA-approved drugs to identify those that effectively inhibited Wnt signaling with minimal off-target effects related to larvae development and survival. We focused on a drug re-purposing/re-positioning screen since the identification of new uses for established drugs leverages the wealth of the pharmacokinetics, pharmacodynamics, and safety data on identified hits. Repurposed drugs progress faster into phases II and III of clinical testing, with a significantly reduced cost [33]. The hits we identified in the zebrafish screen were interrogated for their cross-species effect using complementary human cell reporters. The epidermal growth factor receptor (EGFR) inhibitor Erlotinib emerged as a promising candidate, capable of inhibiting Wnt/β-Catenin signaling in zebrafish and human cells, reducing the expression of Wnt target genes, and blocking the self-renewal of T-ALL stem cells *in vitro* and *in vivo*. In total, these data underscore the potential of zebrafish-based drug screening as a robust strategy for identifying Wnt inhibitors with favorable target specificity and translational relevance.

## 2. Materials and Methods

### 2.1. Zebrafish care and use

Zebrafish care and handling was approved by approved by the University of Kentucky’s Institutional Animal Care and Use Committee, protocol 2019-3399. The *6xTCF/LEF-miniP:dGFP* transgenic zebrafish line [34] and CG1-strain syngeneic zebrafish [35] were used for these studies. Zebrafish were maintained at 28°C with a light/dark cycle of 14:10 hours. Eggs were collected into 1X E3 media (5.0 mM NaCl, 0.17 mM KCl, 0.33 mM CaCl_2_, and 0.33 mM MgSO_4_) containing 200 µL/L of methylene blue.

### 2.2. Cell culture

Human cell lines, including Jurkat T-ALL cells (ATCC, VA, USA, TIB-152) and LS174T colon cancer cells (a gift from Professor Hans Clevers and Marc van de Wetering) were cultured at 37°C in a humidified atmosphere with 5% CO_2_. All media was supplemented with 10% heat-inactivated fetal bovine serum (FBS, Atlanta Biologicals, GA, USA, S11150H). Jurkat cells were cultured in RPMI 1640 (ThermoFisher, MA, USA, 11875119) and LS174T were grown in EMEM (ATCC, VA, USA, 30-2003). The HEK293T cell line containing the TOPFlash reporter has been described previously [36–38]. Mycoplasma testing was routinely performed using the LookOut® Mycoplasma PCR Detection Kit (Sigma, MA, USA, MP0035-1KT) to ensure the absence of contamination before cells were used in experiments.

### 2.3. Drug screening in *6XTCF/LEF-miniP:dGFP* transgenic zebrafish

The *6XTCF/LEF-miniP:dGFP* transgenic zebrafish line was to screen a library of FDA-approved drugs (Enzo Screen-Well® FDA Approved Drug Library V2, NY, USA, version 1.4., BML-2843-0100, Lot No. 06051910A) for modulators of the Wnt/β-catenin/TCF-LEF signaling axis modulators. Screening was carried out as previously described [39, 40]. Briefly, at 24 hours post-fertilization (hpf) zebrafish larvae showing similar GFP fluorescence in the tail fin were selected for use. The larvae were dechorionated and placed individually into a 96-well round bottomed plate with 148.5 µl of 1X E3 media. The drug library was diluted to 100 µM in E3 water immediately before use and drugs were tested at a final concentration of 1 µM by adding 1.5 µl of the 100 µM diluted drug stock to each well. An equivalent volume of DMSO was used as the control. Larvae were incubated for 48hrs at 28°C.

### 2.4. Automated imaging of florescent larvae and fluorescence quantification

GFP fluorescence in the tail fin of the zebrafish larvae was imaged using the Vertebrate Automated Screening Technology (VAST) Bioimager and Large Particle (LP) Sampler, as we have previously described [32]. ImageJ software was used for quantitative analysis by measuring the average area and mean intensity of fluorescence. Clutch-specific fluorescence thresholds were determined using the average GFP fluorescence of DMSO-treated larvae in each clutch.

### 2.5. Caudal fin regeneration assay

To assess the effects of drugs on caudal fin regeneration, zebrafish were subjected to caudal fin amputation, as previously described [39]. Briefly, *6xTCF/LEF-miniP:dGFP* transgenic zebrafish were anesthetized using 100 µl of 4 mg/mL Tricane-S solution in 25 mL of fish system water and placed on a clean 10 cm^2^ petri dish. Using a clean razor blade, the tip of the caudal fin was removed by cutting straight down across the fin. Fish recovered in clean system water for 20 minutes and were then placed into a small tank with 5 µM of drug in 250 mL of system water. Drug was refreshed each day. At 4 days post amputation, fish were euthanized with >400 mg/L Tricane-S solution and fins were imaged. Images were quantified by averaging the longest and shortest point of growth using ImageJ software.

### 2.6. Thawing and propagating zebrafish T-ALL cells

Zebrafish T-ALL with high LIC frequency had been previously described [41]. A frozen viable sample of zebrafish T-ALL #8.1 was thawed for 5 min and then placed on ice. The cells were washed in zebrafish cell media (0.9x PBS + 5% FBS) and then centrifuged at 2500 rcf at 4°C. The supernatant was removed and the pellet re-suspended at 0.5 ml of zebrafish cell media and kept on ice. Approximately 10 µl of the cell suspension was transplanted into the intraperitoneal space of the CG1-strain zebrafish and animals were monitored weekly for leukemia growth.

### 2.7. Limiting dilution transplantation into adult CG1 zebrafish

Fish with ∼80% leukemia burden were used for limiting dilution transplantation to quantify LIC frequency, as previously described [42]. Blood cells were isolated by passing macerated zebrafish through a 40 µM filter and fluorescently labeled leukemia cells were counted. Cell solutions with concentrations of 10, 100 and 1000 leukemia cells per 5 µl were prepared in RPMI 1640 + 10% FBS and treated with 1 µM of drug for 1 hr. Cell viability was confirmed using Trypan blue stain before transplantation. CG1-strain zebrafish were intraperitoneally transplanted at 5 µl of the cell solution/drug mixture. Transplanted zebrafish were moved to 4 L tanks and monitored weekly for leukemia development using a florescent microscope. Data were uploaded to the web-based ELDA (Extreme Limiting Dilution Analysis) statistical software [43] to determine the frequency of self-renewing leukemia cells.

### 2.8. Matrigel Colony formation assay

Actively proliferating Jurkat cells, with > 95% viability as assessed by Trypan blue staining, were treated with drugs at a final concentration of 10 µM and mixed with MethoCult™ H4100 Base medium (Stem cell technologies, Canada, 04100) enriched with 20% FBS. The cell mixture was plated in 6-well plate at a cell density of 2000 cell per well. Plate was imaged after 14 days using the Agilent BioTek Lionheart FX at 4X magnification. Images were stitched using Agilent Gen5 BioTek Microplate Reader and Imager Software and colonies were counted using ImageJ software.

### 2.9. Annexin V-FITC Apoptosis assay

Jurkat cells were treated with drugs at a final concentration of 10µM for 48 hrs. After treatment, cells were collected and stained with the Annexin V-FITC kit (Invitrogen, CA, USA, V13242 A), according to manufacturer’s protocol. Flow cytometry analysis was performed using the BD FACSymphony A3 and analyzed with FlowJo software.

### 2.10. Cell proliferation assay

Jurkat cells were treated with drugs at a final concentration of 10µM for 48 hrs. 100 µL of cells were plated in opaque-walled (white) multi-well plates and 100 µL CellTiter-Glo (Promega, WI, USA, G7572) solution was added avoiding bubbles. Plate was incubated, with shaking for 10 minutes to lyse the cells. Plate was then allowed to stabilize for another 10 minutes at room temperature and luminescence was recorded.

### 2.11. Wnt reporter assay

Wnt reporter assay has been described previously [36–38]. Super 8xTOPFlash (provided by Professor Randall Moon, University of Washington) was sub-cloned into pGL4.83 [*hRlucP*/Puro] and transfected into HEK293T cells to establish a stable cell line containing the TOPFlash reporter. Wnt signaling was activated using 25 mM LiCl or Wnt3A conditioned medium.

### 2.12. Western blot

Cells were lysed in the appropriate volume of lysis buffer containing 50 mM HEPES, 100 mM NaCl, 2 mM EDTA, 1% (v/v) glycerol, 50 mM NaF, 1 mM Na_3_VO_4_, 1% (v/v) Triton X-100, with protease inhibitors. The following antibodies were used: Actin (Sigma, MA, USA, A1978), CyclinB1 (Cell Signaling Technology, MA, USA, 4135), Axin2 (Cell Signaling Technology, MA, USA, 2151), Survivin (Cell Signaling Technology, MA, USA, 2808) and c-Myc (Epitomics, CA, USA, 1472-1).

### 2.13. Seahorse assay

Briefly, 3 x 10^4^ Jurkat cells in 100 µL of medium were seeded in a Cell-Tak (Corning, NY, USA, 354240 coated XF96 Cell Culture microplate and treated with a final concentration of 10 µM Erlotinib and a corresponding amount of the DMSO control in culture media. After 24 hrs., cell culture media were replaced with Seahorse XF modified media. Cells were treated with 1 μM of oligomycin A, 0.6 μM FCCP or mixture of 1 μM of rotenone and 1 μM of antimycin A in standard mitochondrial stress test conditions. For glycolytic stress test, similar plating procedure was followed and 10 mM glucose, 1 μM oligomycin, and 50 mM 2-Deoxy-D-glucose were added to wells.

### 2.14. Statistical analyses

Statistical analyses were carried out using GraphPad Prism. One-way ANOVA with multiple comparisons was done to compare the percent change in GFP fluorescence between drug treated and control. Outliers were determined with the GraphPad Outlier calculator and removed from further analysis. One-way ANOVA with multiple comparisons was also completed to analyze caudal fin growth and colony formation. A Welch’s t-test was used to analyze changes in sphere formation, apoptosis, and proliferation. A multiple Mann-Whitney test was used to analyze difference in mitochondrial and glucose metabolism. A Gehan-Breslow-Wilcoxon test was used to examine the effects of drug on leukemia latency. A P-value of equal to or less than 0.05 was considered statistically significant.

## 3. Results

### 3.1. Validation of *6xTCF/LEF-miniP:dGFP* transgenic zebrafish as a useful tool to identify inhibitors of the Wnt/β-catenin signaling pathway

The Wnt/β-catenin signaling axis is an attractive drug target for cancer, as aberrant Wnt signaling or β-catenin activation is frequently linked with cancer stem cell development and self-renewal. However, Wnt signaling also plays important roles in tissue homeostasis. Drugs targeting this pathway are therefore frequently associated with undesirable side-effects in patients, limiting their clinical utility. There is currently an unmet need for Wnt/β-catenin inhibitors with improved safety profiles. To address this issue, we utilized transgenic strain of zebrafish that harbors a Wnt/β-catenin reporter [39] (*6XTCF/LEF-miniP:dGFP*) in a screen of 773 FDA-approved drugs (**Figure 1A**) to find drugs capable of interfering with the Wnt/β-catenin signaling axis with low toxicity to developing zebrafish larvae. Wnt signaling plays a crucial role in normal zebrafish development and is active in cells within the head and tail-fin regions of 24-hour post-fertilization (hpf) larvae. TCF/LEF transcription factors are the major endpoint mediator of Wnt signaling, and the *6XTCF/LEF-miniP:dGFP* zebrafish line has six TCF/LEF response elements driving transcription of a destabilized GFP [34]. Active Wnt and β-catenin signaling can therefore be visualized in these animals as GFP fluorescence (**Figure 1B**). Imaging of the tail fin, which is relatively thin and 2-D compared to the head and body of the larvae, provides a rapid and accurate means to quantify Wnt signaling responses in an *in vivo* setting [39]. We found the *6XTCF/LEF-miniP:dGFP* animals were sensitive to the known mammalian Wnt pathway modulators XAV939 [44] and BIO [45], resulting in a 90.96% decrease or 79.94% increase in GFP-fluorescence in the tail fin, respectively (**Figure 1C-D**). These data indicate that this model is suitable to identify novel Wnt/β-catenin pathway inhibitors.

**Figure 1:**
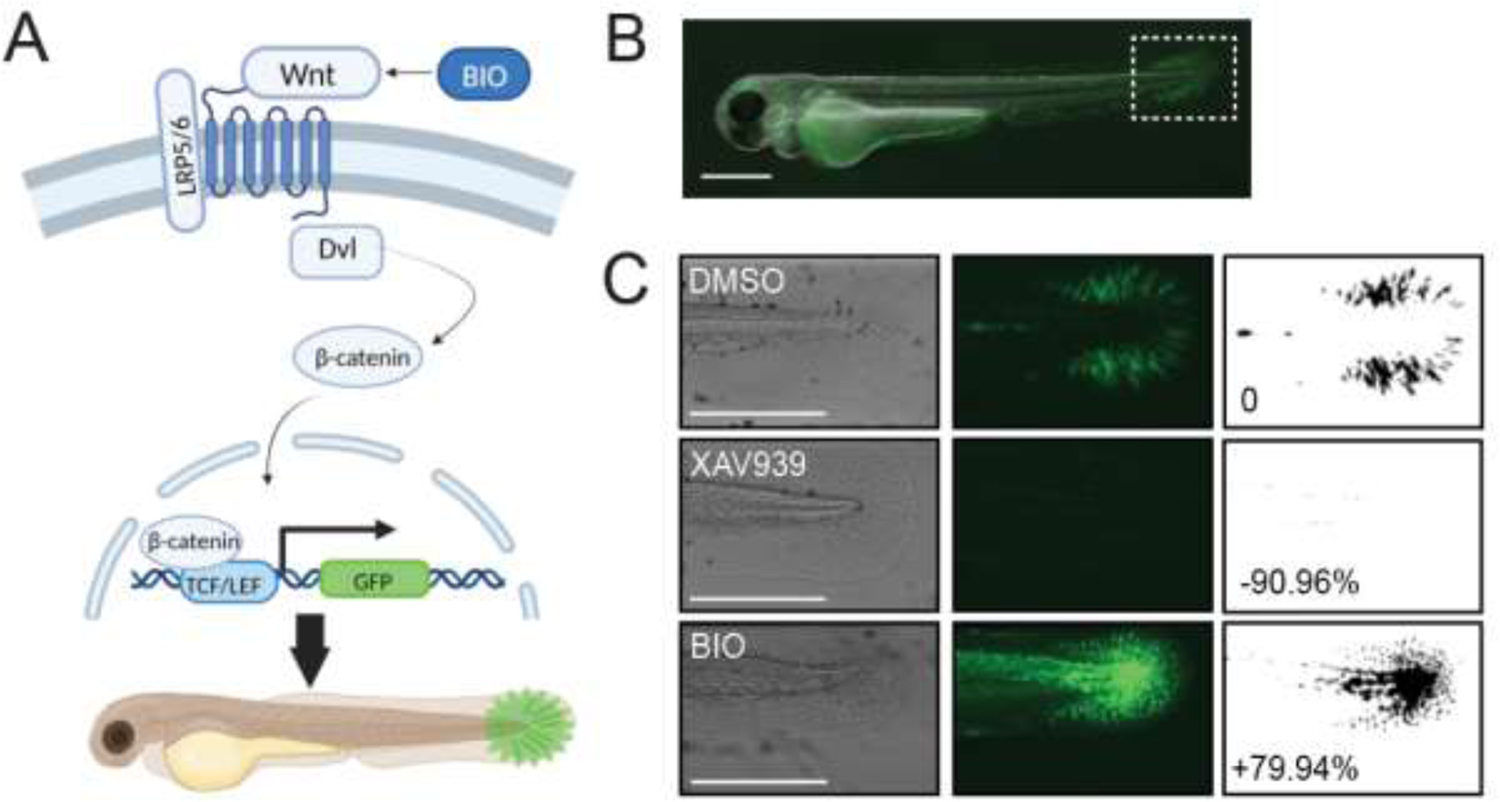
The *6xTCF/LEF-miniP:GFP* zebrafish line produces quantifiable responses to small molecule modulators of the Wnt signaling pathway. (**A**) Schematic of the Wnt/β-catenin GFP reporter the *6xTCF/LEF-miniP:dGFP* transgenic zebrafish line. (**B**) A *6xTCF/LEF-miniP:dGFP* zebrafish larvae at 48 hours post-fertilization (hpf). GFP fluorescence is indicative of active Wnt signaling, and the caudal fin (dashed box) was used for quantification. (**C**) Representative caudal fin fluorescence in *6xTCF/LEF-miniP:dGFP* larvae treated for 24 hr with DMSO, Wnt pathway inhibitor XAV939, or Wnt pathway activator BIO. Panels from left to right show brightfield images, GFP fluorescence, and standardized thresholding of fluorescence using ImageJ software. The percent increase or decrease in fluorescence compared to DMSO is indicated. Scale bars = 500 μm.

### 3.2. Zebrafish provide a rapid and inexpensive method to repurpose FDA-approved drugs as Wnt/β-catenin inhibitors with low *in vivo* toxicity

We applied our drug treatment and imaging workflow to the Enzo Screen-Well FDA-approved drug library. Of the 773 drugs tested, twenty-three resulted in a 95% or greater decrease in GFP-fluorescence *in vivo* compared to the DMSO control (Figure 2A **and Supplemental Table 1**). Upon further validation in larger cohorts of animals, three drugs were fatal to the larvae and were excluded from further study and six of these drugs reduced Wnt/β-catenin signaling, as measured by GFP fluorescence (**Supplemental** Figure 1). Erlotinib and Oxaliplatin, in particular, showed a significant and reproducible inhibition across multiple clutches of *6XTCF/LEF-miniP:dGFP* larvae (Figure 2B-C).

**Figure 2.**
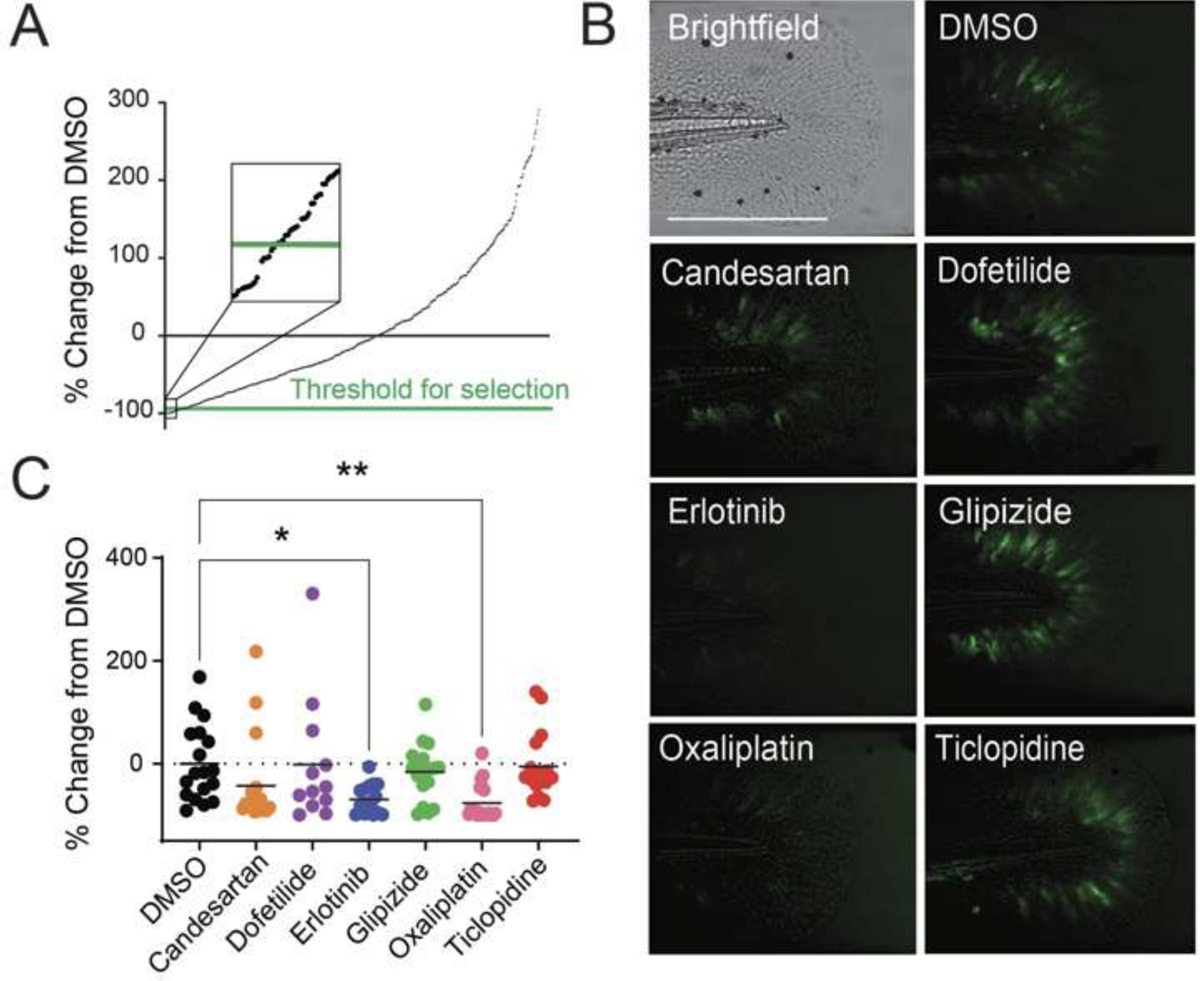
An *in vivo* screen identified FDA-approved drugs with activity against Wnt/β-catenin signaling. (**A**) Quantification of the percent change in GFP fluorescence in *6xTCF/LEF-miniP:GFP* larvae treated with each of the 773 drugs in the Enzo-ScreenWell library, compared to DMSO controls. The threshold for selecting drugs to move forward in the study was a 95% decrease in GFP fluorescence compared to DMSO (inset). (**B**) Representative images of brightfield and GFP fluorescence in the caudal fin of 48 hpf *6xTCF/LEF-miniP:dGFP* transgenic zebrafish treated with DMSO or drugs from the large-scale screen. Scale bar = 500 μm. (**C**) Percent change in GFP fluorescence in larvae treated with the indicated drugs, compared to DMSO, in >10 replicate embryos/drug. * *p* = 0.0327; ** *p* = 0.0095, compared to DMSO.

As an additional validation, we tested whether the six drug hits impacted the ability of adult zebrafish to regenerate their caudal fin after amputation. This regeneration process relies on Wnt signaling [46], and similar assays in zebrafish have been previously used to identify putative Wnt inhibitors [39]. We found that each of our six identified hits significantly reduced caudal fin regeneration by up to 50-90%, and several hits had comparable effects to XAV939, a well-established Wnt pathway inhibitor (Figure 3A-B). A summary of the labeled indications of these drugs and their prior use in cancer research is provided in **Table 1**. To our knowledge, none of the identified hits have been previously linked with the Wnt pathway, suggesting that our *in vivo* screening approach was able to identify novel compounds capable of interfering with the Wnt/β-catenin signaling axis with minimal to no negative effects on zebrafish development and survival.

**Figure 3.**
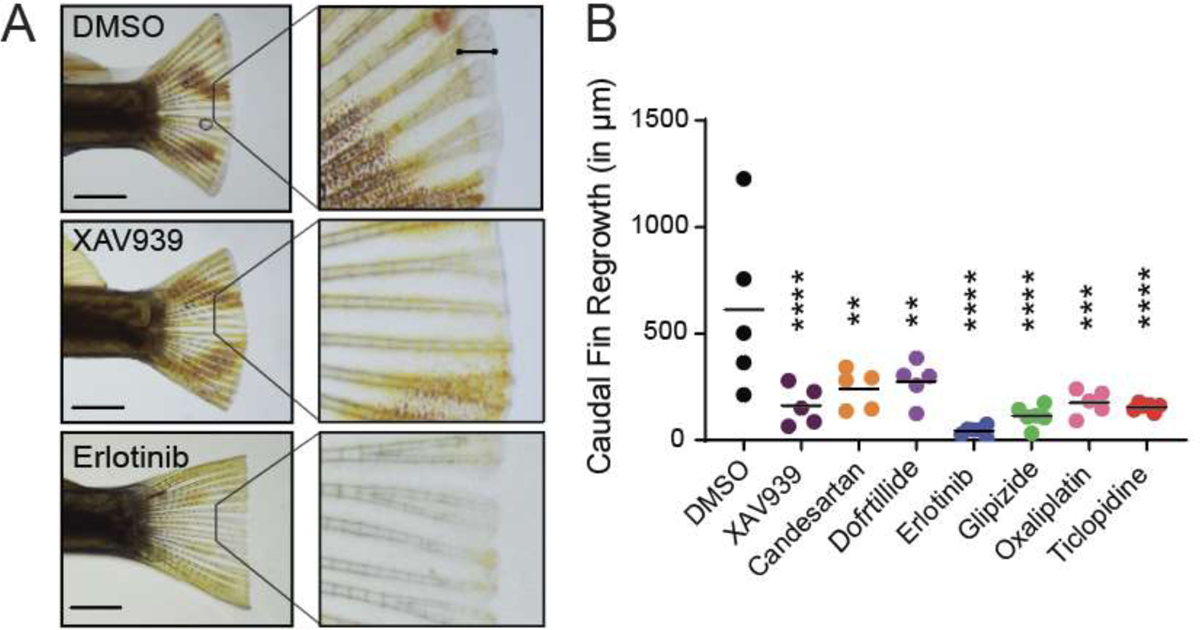
Drugs identified in the larval zebrafish screen also inhibit Wnt-dependent caudal fin regeneration in adult zebrafish. (**A**) Representative images of zebrafish 4 days post-amputation of the caudal fin. The area of regrowth is non-pigmented, denoted in the upper right box by the black line. Animals were treated at the time of amputation with DMSO, 1 μM of the Wnt inhibitor XAV939 or 5 μM Erlotinib. Scale bar = 2 mm. (**B**) Quantification of the length of fin regrowth 4 days post-amputation. Each data point represents one animal treated with the xx μM of the drug indicated at the time of amputation. ** *p* ≤ 0.01; *** *p* =0.0002; **** *p* ≤ 0.0001, compared to DMSO treatment.

**Table 1:**
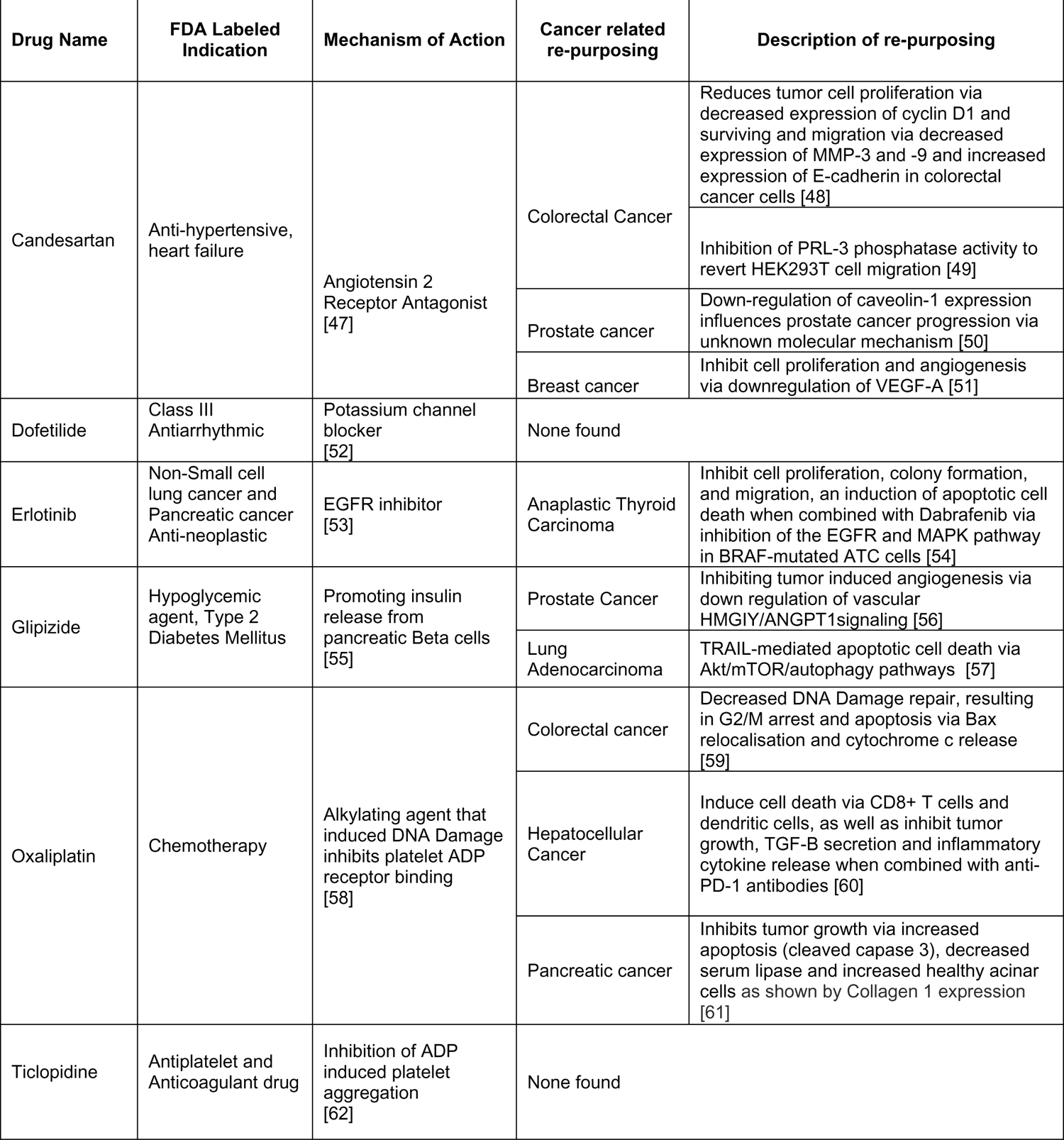
Summary of the FDA-approved drug hits with capability of inhibiting Wnt/β-catenin signaling in human and zebrafish cells.

### 3.3. Erlotinib inhibits the Wnt/β-Catenin signaling axis across species

To determine if the inhibitory effects of the drugs are conserved from zebrafish to human, we tested each drug in a TOPFlash luciferase reporter assay, which utilizes a human HEK293T cell line harboring a TCF/LEF responsive element that drives expression of luciferase when Wnt/β-catenin signaling is activated [36–38]. The cells were treated with either Wnt3a or Lithium Chloride (LiCl) to narrow where in the Wnt signaling pathway our drugs of interest might act. Wnt3a activates Wnt signaling at the Frizzled cell surface receptor [63], while Lithium Chloride (LiCl) inhibits GSK3 and the destruction complex, leading to β-catenin stabilization and downstream transcription of Wnt target genes [64] (Figure 4A), Among the six drugs, Erlotinib significantly inhibited both the luciferase activity induced by Wnt3A (Figure 4B) and LiCl (Figure 4C). In LS174T colorectal cancer cells, which have harbored a mutated form of β-catenin that is resistant to degradation, Erlotinib treatment reduced expression of the endogenous Wnt target genes, c-Myc, Axin-2, Survivn (Figure 4D). Together, these data indicate that Erlotinib acts downstream of the destruction complex to inhibit transcript of β-catenin target genes in both zebrafish and human cells.

**Figure 4.**
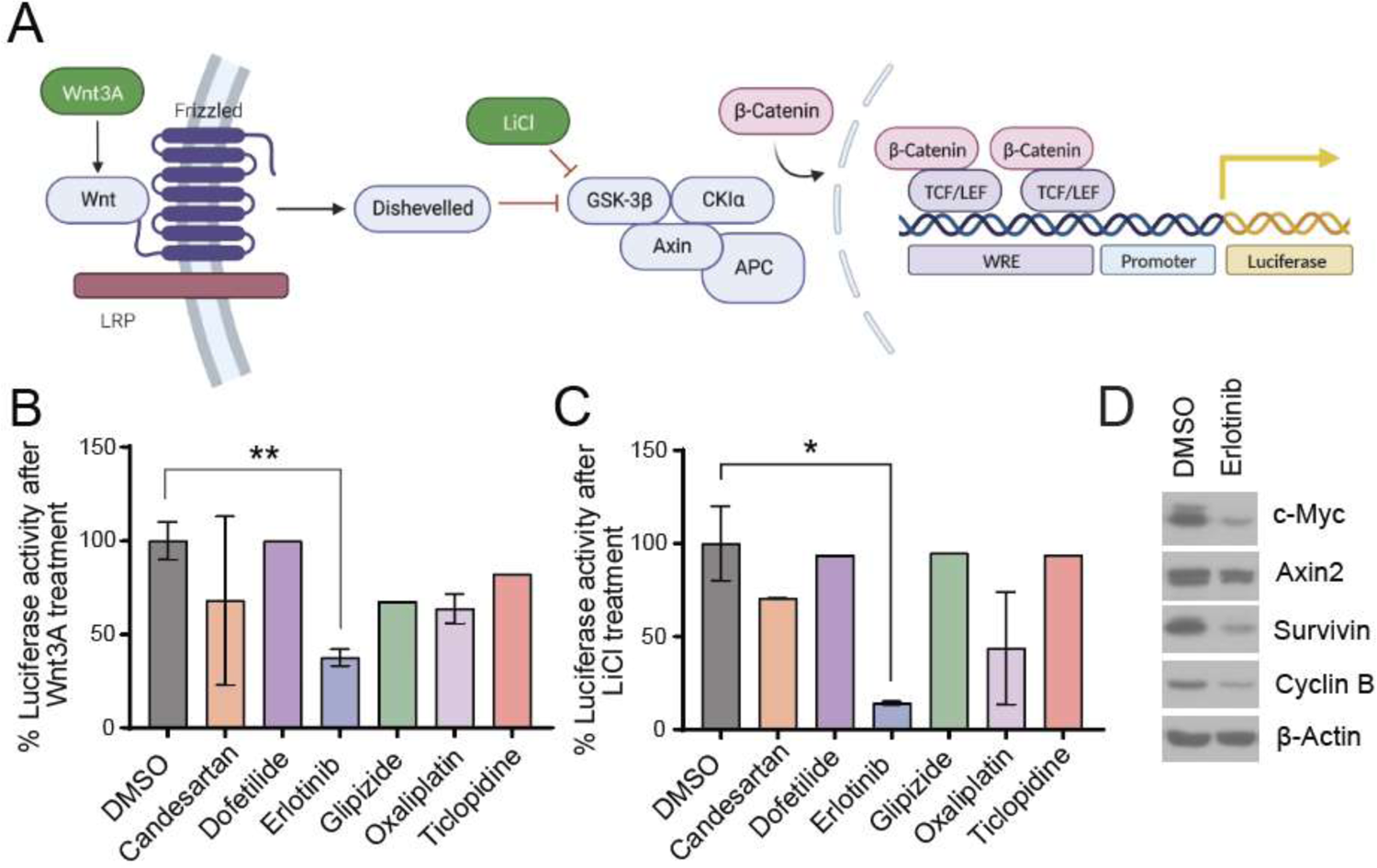
Erlotinib treatment reduced the expression of Wnt/*β-Catenin* target genes in human cells. (**A**) A schematic of the TOPFlash luciferase assay to assess Wnt/β-catenin activity in human embryonic kidney (HEK293T) cells. Wnt3A is a general Wnt activator and LiCl is an GSK3 inhibitor that results in the stabilization of β-catenin. TOPFlash reporter cells were treated with conditioned media containing Wnt3a (**B**) or 25 mM LiCl (**C**) and DMSO or the indicated drugs for 24 hr. The average luciferase activity in the DMSO-treated cells was considered 100% activity, and data from individual DMSO or drug-treated wells were compared to this value. DMSO, Candesartan, Erlotinib, and Oxaliplatin were tested in 3 individual assays, the remaining drugs were used in the assay one time. * *p* = 0.011; ** *p* = 0.0041 (**D**) Western blot of LS174T colorectal cancer cell lysates treated with DMSO or 10 µM Erlotinib for 24 hr. Blots were probed with the indicated antibodies for Wnt target genes, β actin was used as a loading control.

### 3.4. Erlotinib prevents self-renewal of leukemia initiating cells *in vitro*

Self-renewal of T-cell Acute Lymphoblastic Leukemia is maintained at least in part through β-catenin signaling [20]. Leukemia stem cells, also known as leukemia initiating cells (LICs), are thought to drive relapse formation in T-ALL, and the inability of current cytotoxic chemotherapy strategies to reliably target LICs remains a major clinical issue for this malignancy [65]. We treated a human T-ALL cell line with Erlotinib to determine the effect of the drug on LIC self-renewal using an *in vitro* sphere formation assay.

Formation of spherical aggregates in Matrigel is a key feature of leukemia initiating cells [66] and sphere formation assays are commonly used to evaluate the impact of various experimental conditions on self-renewal in a variety of cancers [67–69]. We found that Erlotinib treatment reduced the number of spheres formed by 45% compared to DMSO (*p* = 0.0076, Figure 5A and B**, Supplemental** Figure 2). There was no significant difference in the size of the spheres that did form in the Erlotinib treated groups, suggesting the effect of the drug is likely due to an impact on self-renewal rather than proliferation rates (Figure 5C). Erlotinib treatment induced a significant ∼4-fold increase in apoptosis in T-ALL cells after 24 hr (*p* = 0.0097, Figure 5D), but the overall cell number was only slightly reduced at later timepoints (*p* = 0.1913, Figure 5E). We next examined cellular bioenergetics, as some Wnt pathway inhibitors have recently been found to act via reducing ATP synthesis [37]. However, we found T-ALL cells had no significant change in ATP production after Erlotinib treatment (Figure 4E). Mitochondrial respiration and the overall glycolytic capacity of Erlotinib treated T-ALL cells was slightly altered (Figures 4E-F). The reasons for these changes are unknown, although glycolysis has been previously linked to LIC function[70]. Taken together, these data suggest that the major functional effect of Erlotinib in T-ALL cells is on reducing LIC self-renewal, rather than overall cell survival.

**Figure 5.**
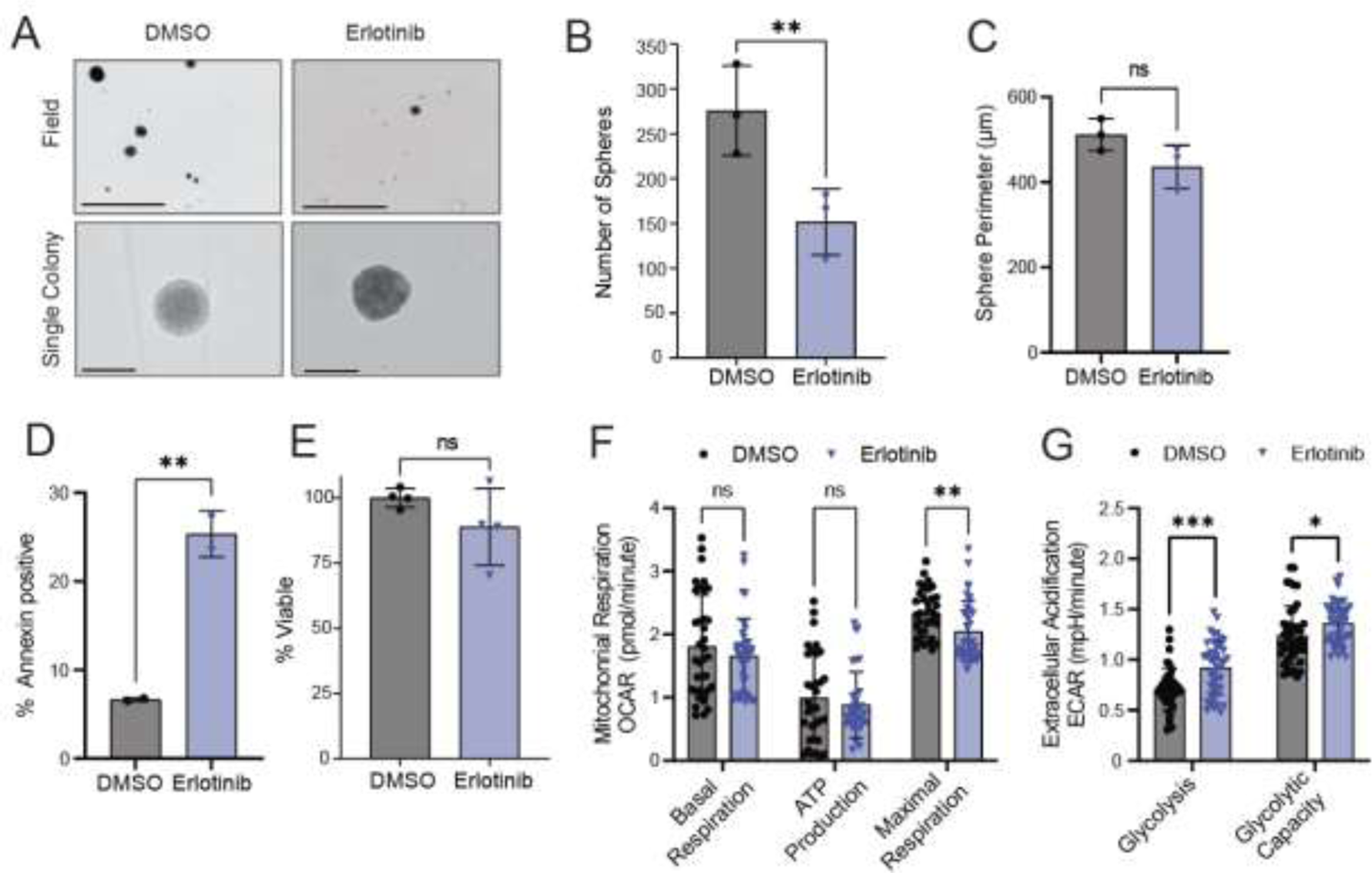
Erlotinib reduces self-renewal of T-ALL cells *in vitro*. (**A**) The Jurkat T-ALL line was treated with DMSO or 10 μM Erlotinib and used in a sphere formation assay. Representative images of the field view at 4x magnification (top, scale bar is 1000 μm) and individual colonies (bottom, scale bar is 100 μm) are shown after 14 d of growth. (**B**) Quantification of the number of colonies per well, from (A). *\*\* p* = 0.0076 compared to DMSO treatment. (**C**) Measurement of the perimeter of the colonies identified in (B). (**D**) Percent of cells that were Annexin V positive after 24 hr treatment with DMSO or 10 μM Erlotinib, ** *p* = 0.0097). (**E**) Percent change in the viability of cells after 48 hr of the indicated treatment, using Cell Titer Glo to quantify metabolically active cells. (**F**) The normalized oxygen consumption rate following a mitochondrial stress test and (**G**) normalized extracellular acidification rate following a glycolytic stress test of T-ALL cells treated with DMSO or 10 μM Erlotinib. * *p* = 0.034; ** *p* = 0.0057; *** *p* = 0.00021, compared to DMSO. For all, *ns* = not significant and each data point is an individual well from experimental replicates.

**Figure 6.**
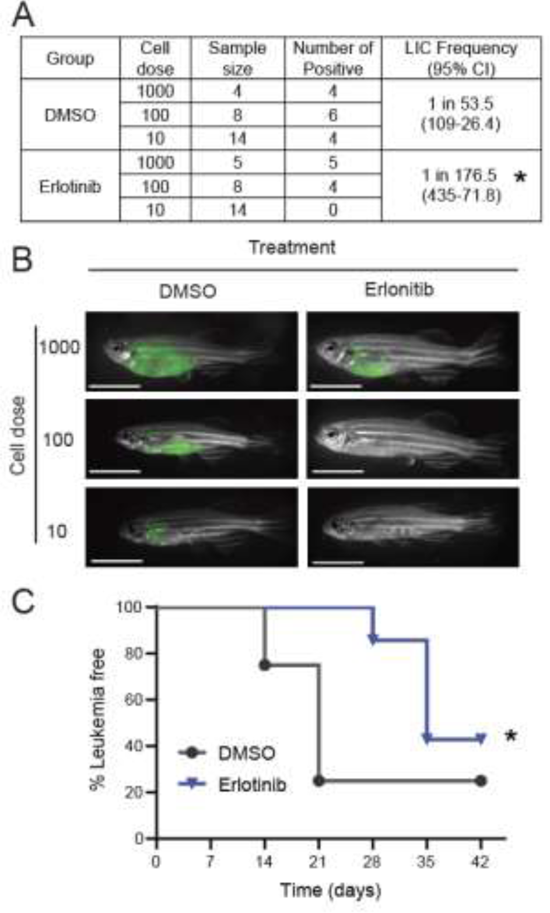
Erlotinib treatment reduced LIC frequency in an *in vivo* T-ALL model. (**A**) Data from a limiting dilution transplantation assay using a GFP-expressing zebrafish T-ALL, showing the number of leukemia-positive animals after transplant with increasing doses of cells, treated with either DMSO or 1 μM Erlotinib. The calculated LIC frequency and 95% confidence interval is shown, \**p =* 0.0352. (**B**) Representative images of animals following 28 days of engraftment, scale bar is 10 mm. (**C**) Survival curve of animals transplanted with 100 cells, data shown are the time required for > XX % of the animal to become GFP-positive, representing significant leukemia burden. * *p =* 0.0384 compared to DMSO treatment.

### 3.5. Erlotinib reduced frequency of T-ALL leukemia initiating cells *in vivo*

We next utilized a zebrafish Myc-induced T-ALL model to determine the effect of Erlotinib on T-ALL progression and self-renewal *in vivo*. A previously isolated zebrafish T-ALL with a relatively high rate of self-renewal [41] was treated with 1 μM Erlotinib or an equal volume of DMSO and transplanted into syngeneic adult recipient zebrafish at limiting dilution, as previously described [42]. Erlotinib caused a three-fold decrease in LIC frequency compared to DMSO control (*p*=0.0352, Figure 5A). Leukemias that did arise also grew significantly slower in the Erlotinib treated group, with animals requiring 28 days for leukemia reformation after transplant, compared to 14 days in DMSO treated controls (*p* = 0.0384, Figures 5B-C). In total, these data demonstrate that Erlotinib has activity against LICs in T-ALL in an *in vivo* system.

## 4. Discussion

Dysregulation of the Wnt/β-catenin signaling pathway has been implicated in the progression and maintenance of many different types of cancer, but the clinical development of Wnt pathway inhibitors has faced challenges. Wnt signaling plays a central role in development, tissue homeostasis and stem cell maintenance [71], so drugs that target this pathway often induce undesirable side-effects related to tissue regeneration, such as bone metabolism disorders, diarrhea, vomiting and others. Such toxicities might not be completely reversible and limit the current clinical utility of Wnt/β-catenin pathway inhibitors [72]. We used a zebrafish drug screening platform to identify FDA-approved drugs that can inhibit Wnt/β-catenin signaling with minimal side-effects in normal larval development. We showed that compounds identified from this screen are also effective in human cells, and demonstrated the ability of a top hit, the EGFR inhibitor Erlotinib, to inhibit self-renewal of leukemia initiating cells in T-ALL, which have previously been determined to have high β-catenin activity [19].

While drug repurposing screens have been employed previously in efforts to identify modulators of Wnt/β-catenin signaling, to our knowledge, our study is the first to use zebrafish models for this particular purpose at a large scale. Typical *in vitro* approaches are anchored in high-throughput screens using cell-based assays with Wnt-responsive reporters. These screens have identified FDA approved drugs with potential anti-Wnt properties, such as nonsteroidal anti-inflammatory drugs (NSAIDS) and statins, which are still undergoing pre-clinical and clinical testing [73–75]. However, cell-based assays lack the complexity of a whole organism, potentially leading to false-positives or overlooking compounds that require systemic interactions for their full effect. The zebrafish TCF/LEF reporter line allowed us to visualize the effects of compounds on Wnt/β-catenin signaling activity in real time, while simultaneously screening for toxicities often related to Wnt inhibition, such as developmental defects. The hits that we identified in zebrafish, and subsequently validated in human cells, have not previously been found using *in vitro* approaches and expand the pool of drugs that could be repurposed to target Wnt/β-catenin signaling in the cancer clinic. Our data demonstrate that zebrafish models can bridge the gap between *in vitro* assays for Wnt/β-catenin inhibition and costly and time-consuming *in vivo* mouse models by combining the many of the advantages of high-throughput screening with the physiological relevance on an intact organism, which may ultimately propel drug discovery related to the Wnt/β-catenin signaling pathway into newer, more promising horizons.

We identified Erlotinib as having potent inhibitory activity against the Wnt/β-catenin signaling pathway in both zebrafish and human cells. Erlotinib, marketed under the brand name Tarceva, is currently used in the oncology clinic as a tyrosine kinase inhibitor, primarily targets the Epidermal Growth Factor Receptor, and has been a front-line treatment for patients with certain types of lung and pancreatic cancers for nearly twenty years [76]. Erlotinib has previously been found to reduce β-catenin expression in squamous cell carcinoma cell lines [77] and to decrease S675 phosphorylated β-catenin in endometrial cancer cell lines, causing β-catenin to be destabilized and degraded [78]. The mechanisms by which Erlotinib might target Wnt/β-catenin signaling are likely indirect, and a growing body of evidence suggests there is cross-talk between EGFR and Wnt signaling. Components of the EGFR pathway translocate into the nucleus and promote β-catenin’s transcriptional activity [79, 80]. EGFR can also activate PI3K/Akt signaling pathway, which, in turn, promotes Wnt signaling [81, 82]. Erlotinib’s on-target inhibition of EGFR could result in decreased Wnt/β-catenin signaling in either of these cases. While cell signaling is context-dependent, and often varies among cell types and conditions, we found Erlotinib to be an effective Wnt/β-catenin inhibitor in developing zebrafish, human embryonic kidney cells, and human and zebrafish T-ALL. This provides some confidence in the drug action, although the precise mechanism of inhibition remains to be determined.

We found that Erlotinib could significantly reduce the self-renewal capabilities of LICs in both human and zebrafish T-ALL. This observation aligns well with the established role of β-catenin signaling in LIC function in T-ALL [83]. Erlotinib has been found to block cancer stem cell function in other cancer types as well. It significantly reduced the colony forming ability of colorectal cancer cells [84], which also rely on β-catenin signaling for stem cell function. In a lung cancer xenograft model, Erlotinib showed selective activity against lung cancer stem cells compared to their more differentiated counterparts, resulting in reduced tumor aggressiveness [85]. Erlotinib was also found to reduce glioblastoma stem cells [86] and push head and neck squamous cell carcinoma stem cells into a more differentiated state [87]. The ultimate translational impact of these findings, including our own, will rely on further validation in mammalian systems or patient-derived xenografts. However, Erlotinib’s FDA approval and well-documented safety profile [88] may provide an exciting opportunity for its use in combination with conventional chemotherapies to target cancer stem cells in a variety of cancer types, including T-ALL.

## 5. Conclusion

Our study employed a zebrafish-based drug screen and identified the FDA-approved drug Erlotinib as a promising compound for targeting the Wnt/β-catenin signaling pathway. This pathway plays a crucial role in embryonic development and tissue repair but current inhibitors have side effects that limit clinical utility. By using a TCF/LEF transgenic zebrafish model, we efficiently screened over 700 FDA-approved drugs and identified Erlotinib as a hit compound that effectively inhibits the transcription of Wnt/β-catenin target genes in zebrafish and human cells. Our findings highlight the potential of Erlotinib as a therapeutic strategy for targeting self-renewal of leukemia initiating cells in T-ALL, opening possibilities for further validation and its combination with conventional therapy to improve patient outcomes.

## 6. List of abbreviations

Abbreviation: Definition

ADP: Adenosine diphosphate

AMP: Adenosine monophosphate

AMPK: AMP-Activated Protein Kinase

ANOVA: Analysis Of Variance

APC: Adenomatous polyposis coli

ATP: Adenosine triphosphate

CSC: Cancer Stem Cell

DMSO: Dimethyl sulfoxide

Dvl: Dishevelled Protein

EDTA: Ethylenediaminetetraacetic Acid

EGFR: Epidermal Growth Factor Receptor

ELDA: Extreme Limiting Dilution Analysis

EMEM: Eagle’s Minimum Essential Medium

ETC: Electron Transport Chain

FACS: Fluorescence-Activated Cell Sorting

FBS: Fetal Bovine Serum

FDA: Food and Drug Administration

FITC: Fluorescein Isothiocyanate

Fzd: Frizzled Receptor Protein

GFP: Green Fluorescent Protein

GSK3β: Glycogen synthase kinase-3 beta

HEPES: N-2-Hydroxyethylpiperazine-N-2-Ethane Sulfonic Acid

Hpf: hours post-fertilization

IACUC: Institutional Animal Care and Use Committee

LDA: Limiting Dilution Assay

LEF: Lymphoid Enhancer Factor

LIC: Leukemia Initiating Cells

LiCL: Lithium Chloride

LP: Sampler Large Particle Sampler

MRD: Minimal Residual Disease

PBS: Phosphate-Buffered Saline

T-ALL: T-cell Acute Lymphoblastic Leukemia

TCF: T-cell Factor

VAST: Vertebrate Automated Screening Technology

Wnt: Wingless-Related Integration Site

## 7. Data Availability

All original data are included in the article and supplementary material. Further inquiries can be directed to the corresponding author.

## 8. CRediT author statement

Author contributions are as follows: conceptualization, MAA, MGH, and JSB; methodology, MAA, AHC, and MGH; formal analysis, MAA; investigation, MAA, AHC, MGH, WZ, ECA, MW, and SS; resources, CL and JSB; writing (original draft), MAA and AHC; writing (review and editing), JSB; visualization, MAA and AHC; supervision, CL and JSB; funding acquisition, JSB.

