## Supplemental Figures for "Zebrafish Drug Screening Identifies Erlotinib as an Inhibitor of Wnt/β-Catenin Signaling and Self-Renewal in T-cell Acute Lymphoblastic Leukemia"

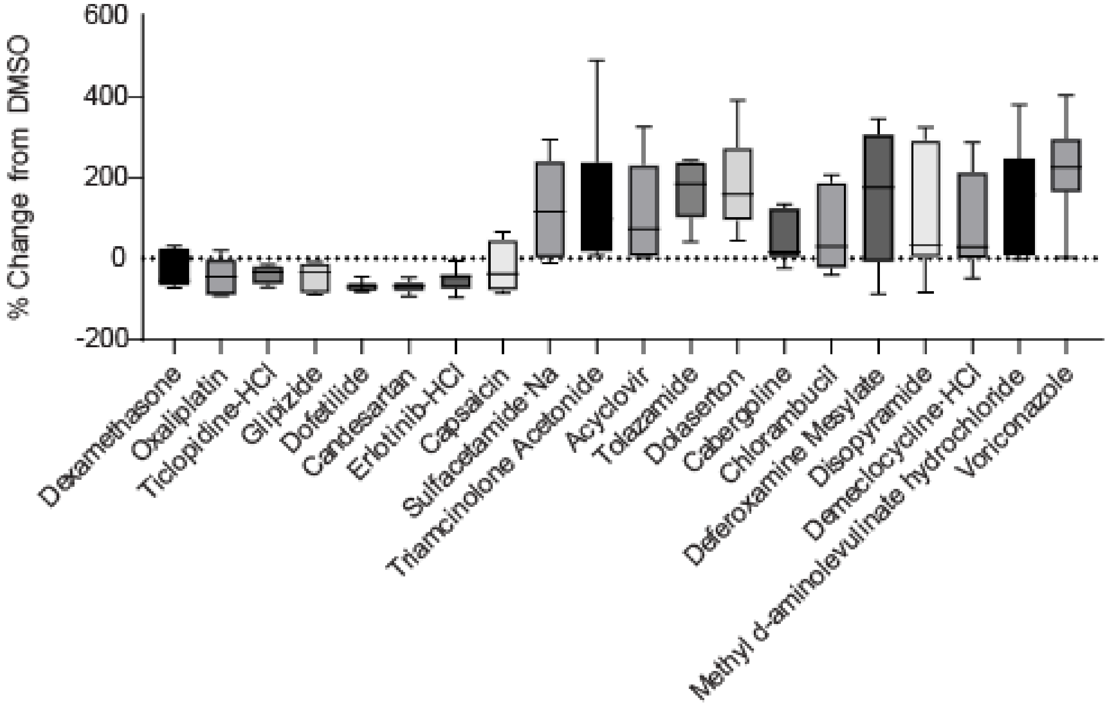


**Supplemental Figure 1. Secondary validation of the compounds identified 6 hit compounds.** The indicated compounds were used to treat 24 hour post-fertilization *6xTCF/LEF-miniP:dGFP* zebrafish, at eight animals per group. The percent chance in fluorescence intensity, which an indication of Wnt signaling activity, was analyzed as described and shown in comparison to DMSO.

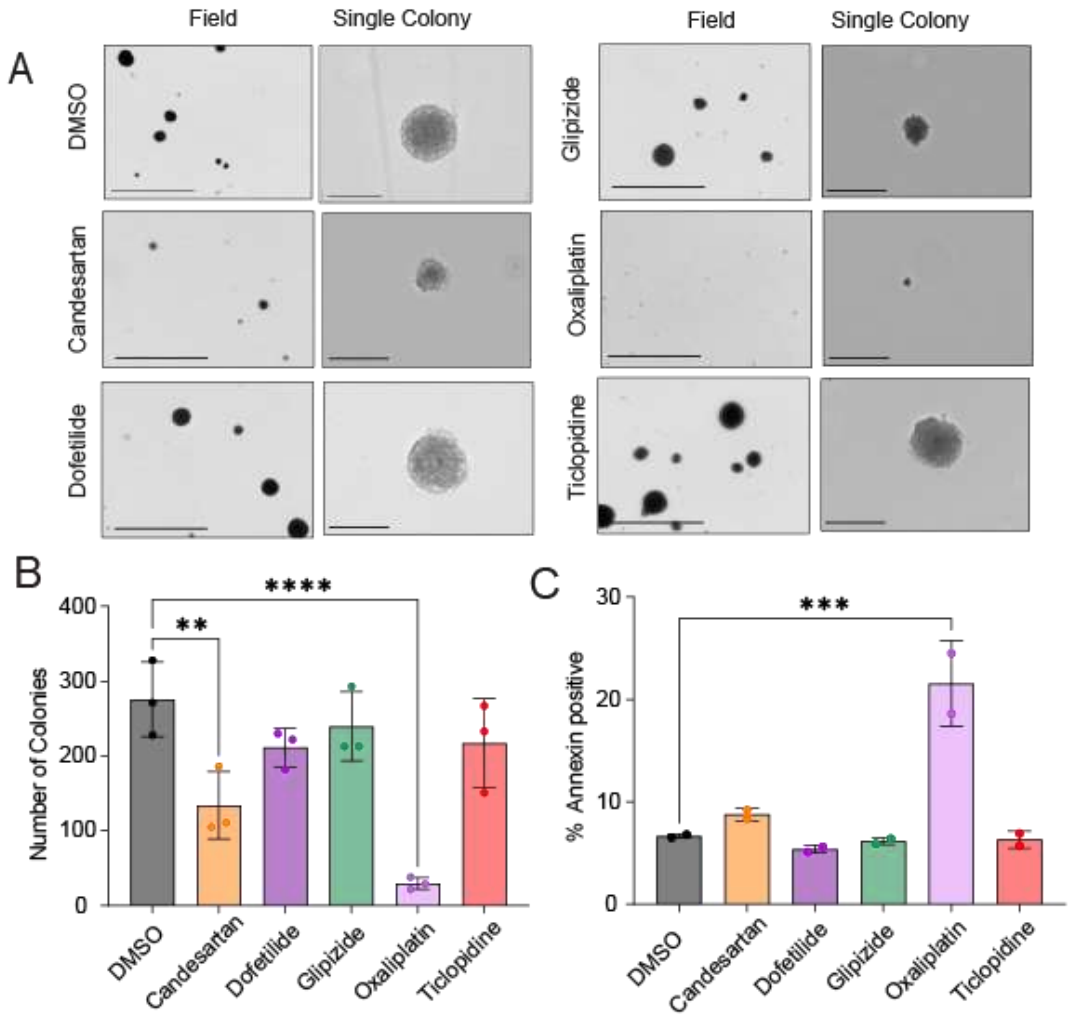


**Supplemental Figure 2. Sphere formation assay and Annexin V staining for the rest of the drug resulting from the screen.** (**A**) The Jurkat T-ALL cell line was treated with DMSO or 10 μM of the indicated drugs and used in a sphere formation assay. Representative images of the field view at 4x magnification (scale bar is 1000 μm) and individual colonies (scale bar is 100 μm) are shown after 14 d of growth. (**B**) Quantification of the number of colonies per well, from (A). ** $p$ = 0.0025 and **** $p \le 0.0001$compared to DMSO treatment. (**C**) Percent of cells that were Annexin V positive after 24 hr treatment with DMSO or 10 μM of the described drugs, ** $p$ = 0.0006).
